# Hitting a moving target: Microbial evolutionary strategies in a dynamic ocean

**DOI:** 10.1101/637272

**Authors:** Nathan G. Walworth, Emily J. Zakem, John P. Dunne, Sinéad Collins, Naomi M. Levine

## Abstract

Marine microbes form the base of ocean food webs and drive ocean biogeochemical cycling. Yet little is known about how microbial populations will evolve due to global change-driven shifts in ocean dynamics. Understanding adaptive timescales is critical where long-term trends (e.g. warming) are coupled to shorter-term advection dynamics that move organisms rapidly between ecoregions. Here we investigated the interplay between physical and biological timescales using a model of adaptation and an eddy-resolving ocean circulation climate model. Two criteria (α and β) were identified that relate physical and biological timescales and determine the timing and nature of adaptation. Genetic adaptation was impeded in highly variable regimes (α<1) but promoted in more stable environments (α>1). An evolutionary trade-off emerged where greater short-term transgenerational effects (low-β-strategy) enabled rapid responses to environmental fluctuations but delayed genetic adaptation, while fewer short-term transgenerational effects (high-β-strategy) allowed faster genetic adaptation but inhibited short-term responses. Our results suggest that organisms with faster growth rates are better positioned to adapt to rapidly changing ocean conditions and that more variable environments will favor a bet-hedging, low-β-strategy. Understanding the relationship between evolutionary and physical timescales is critical for robust predictions of future microbial dynamics.

## Introduction

Ocean circulation advects particles rapidly throughout the ocean basins^1-3^ resulting in variability in physical and chemical properties along Lagrangian trajectories. This variability is both predictable (e.g. diurnal and seasonal cycles) and stochastic (e.g. mesoscale eddies) and is overlain on top of longer-term trends such as those driven by climate change. Understanding the interplay between physical variability in the ocean and the timescales of biological responses to this variability is critical for accurately predicting future shifts in microbial diversity, ecosystem dynamics, and biogeochemical cycling.

Microbial populations – defined as clusters of closely related organisms exhibiting population-specific gene flow – are acted upon by both natural selection and neutral evolutionary processes. While previous work has suggested that marine microbes can evolve faster through neutral genetic processes than they can be dispersed in ocean currents^2^, we have little understanding of how adaptation of microbial populations to different ecological niches through natural selection^4,5,6,7^ interacts with physical timescales in the oceans. Experimental evolution studies have demonstrated relatively fast timescales (<350 generations) of selective adaptation for marine microbes under constant conditions^8^, and suggested that fluctuations may impact the outcome of evolution^9^, consistent with theory^10^. However, our understanding of how marine microbial evolution will proceed *in situ* in a fluctuating environment remains in its infancy.

Heritable variation in fitness can be generated through a range of processes from transgenerational plasticity to genetic mutations. These processes for generating and transmitting trait variation can be classified on a spectrum from *fast variation, low transmission* (LT) to *slow variation, high transmission* (HT) modifications^11^. HT modifications are relatively rare but have a high probability of being transmitted to offspring through a large number of cell divisions. Classic examples of HT modifications are point mutations, genome rearrangement, horizontal gene transfer, and transposon insertions. In contrast, LT modifications are common relative to HT modifications but have a lower probability of being transmitted to offspring. LT modifications include – but aren’t limited to – transgenerational plastic effects, and changes to DNA methylation and acetylation patterns (i.e. epigenetics). Theoretical^12,13^ and empirical^15^ data suggest that HT and LT modifications acting together best explain patterns of microbial evolution on timescales of hundreds of generations.

Immediately following environmental change, LT modifications may allow for flexible and rapid exploration of phenotypic space. This can result in enhanced rates of adaptation to the new environment (increase in fitness) relative to what would be expected due to HT modifications alone^13-16^. However, once a perturbation is removed, the fitness benefits from LT modifications will be lost from the population more quickly than would be expected from HT modifications alone. Currently, we lack an understanding of how the interplay between LT and HT modifications can affect both the timescale and outcome of marine microbial adaptation in a dynamic physical environment. Here we use two numerical models to link adaptive and physical timescales. We show how the degree of environmental stability along Lagrangian trajectories relates to evolutionary tipping points where more stable trajectories favor evolution driven by HT modifications while variable trajectories favor strategies based on LT (i.e. non-genetic) modifications. Understanding the trade-offs between different evolutionary strategies in the context of a fluctuating environment will allow for improved predictions of how general patterns of trait distributions^18^ among marine microbial functional groups^19^ might shift in a changing world.

## Variable Selection Pressures

When considering adaptation in a variable environment, it is necessary to clearly define the effects of selection pressure across different types of environments. We distinguish between two types of environments: the ‘new’ environment where populations are under directional selection (the selective fixation of new beneficial alleles where the population is in the process of adapting); and the ‘ancestral’ environment where the population is well adapted and under stabilizing selection (i.e. the selective removal of new alleles, which are deleterious). We investigated the interactions between evolutionary timescales (changes in population fitness) and physical timescales (timescales of environmental fluctuations), using an individual based model of adaptation modified from Fisher’s model^13,20^. In the model simulations, the population moved between the ‘new’ and ‘ancestral’ environment with varying frequencies. Adaptation -- increases in fitness in the ‘new environment’ -- was driven by both LT and HT modifications. Critically, LT modifications were introduced at a higher frequency than HT modifications but were also associated with a transmission timescale or reversion rate (*Methods*). As a result, the model captured both the high frequency occurrence of LT modifications (e.g. transgenerational plastic responses) in populations following an environmental change and the degradation of this signal over several generations once the environmental cue was removed. In contrast, HT modifications (e.g. genetic mutations) occurred at low frequencies in the population, but were transmitted with high fidelity between generations.

Three critical timescales emerged in the model (*Figure 1*); 1) the time spent in each environment measured in generations (τ_f_), 2) the emergent timescale required for a beneficial HT modification to fix in the population through a selective sweep^5^ once it occurred in an individual (τ_HT_), and 3) the specified transmission timescale for LT modifications (τ_LT_). The model was run using a range of τ_f_ and τ_LT_ (*Methods*). τ_HT_ is an emergent property of the model that varied as a function of HT modification supply and effect (*Figure S1*). The strength of population size and stabilizing selection were also tested and shown to not change overall patterns (*Supplement S1&S2*).

**Figure 1:**
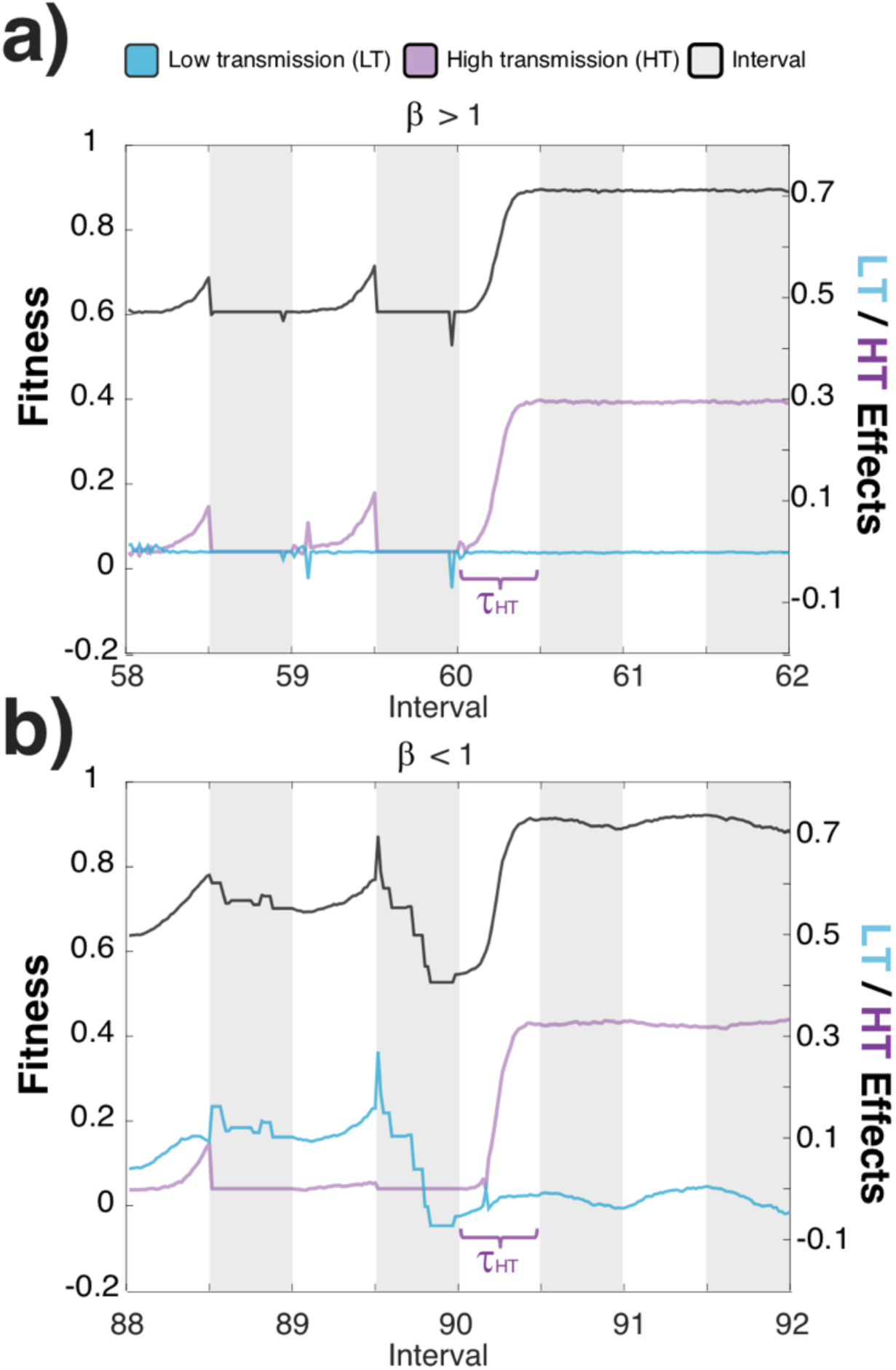
Illustrative example of model dynamics for a high-β (a) and low-β (b) simulation. Fitness changes (black line) are primarily driven by HT modifications (purple line) in the high-β simulation and by both HT and LT (blue line) modifications in the low-β simulation. The time-to-sweep (τ_sweep_) is longer for the low-β simulation (b) than the high-β simulation (a). White shading denotes the ‘new environment’ while grey shading denotes the ‘ancestral environment’.

In all model simulations, fitness increased rapidly with exposure to the ‘new’ environment, consistent with laboratory experiments^21-23^. With stabilizing selection applied during the ‘ancestral environment’ periods, selective sweeps driven by HT modifications emerged if the fluctuation intervals (τ_f_) were long enough. We identified two dimensionless criteria that determined model behavior across the wide range of parameter values tested:

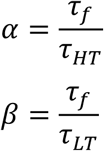

When α <1, the timescales of environmental variability (τ_f_) were short relative to the fixation timescale for HT modifications (τ_HT_) and so selective sweeps based on HT modifications were inhibited (*Figure 2a*). Conversely, when α >1, HT selective sweeps always occurred and the time to sweep (τ_sweep_) decreased as τ_f_ increased. In other words, longer exposure times to a new environment drove higher rates of genetic adaptation to that environment, consistent with models using genetic mutation only^24^.

**Figure 2:**
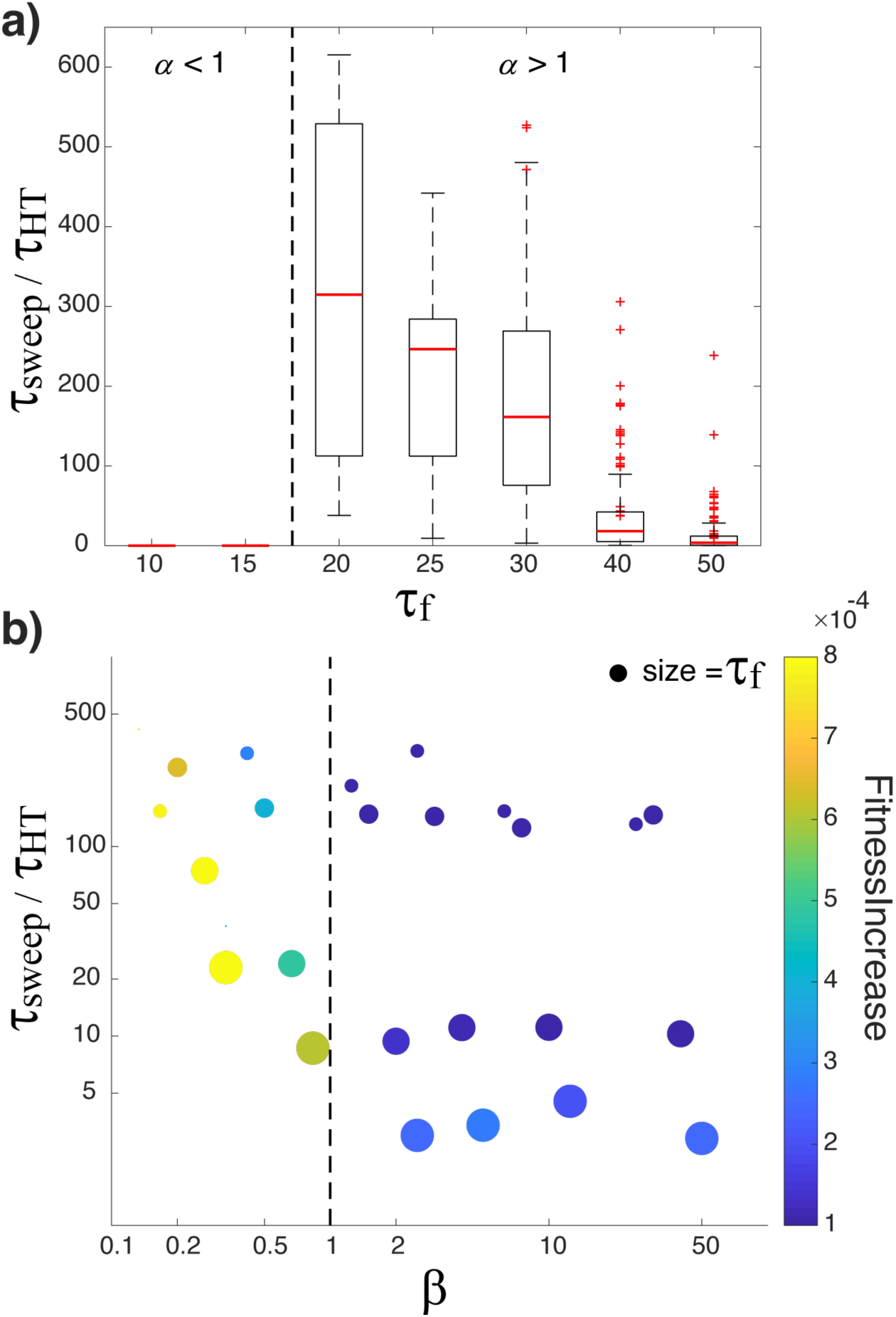
Timescales and outcomes of adaptation are determined by the values α and β. Panel a illustrates the α criteria by showing the impact of environmental fluctuations (τ_f_) on τ_sweep_ normalized to τ_HT_. Panel b illustrates the trade-off associated with a low-β strategy by showing the relationship between the rate of fitness increase in a new environment (colorbar) with τ_sweep_ normalized to τ_HT_. In panel b, τ_f_ is represented by the size of the symbol.

The second criteria, β, identifies a key evolutionary trade-off for organisms in a fluctuating environment. When β>1, HT modifications drove adaptive fitness changes while LT modifications played a minor role, resulting in little or no short-term responses (i.e. fitness changes) to environmental fluctuations (*Figure 1a*). However, when β<1, LT modifications enabled short-term fitness responses to environmental fluctuations both before and after a HT selective sweep (*Figure 1b*). Although simulations with β<1 had a more rapid response to environmental change (faster increase in fitness), it also took longer for a HT sweep to occur (larger τ_sweep_) than simulations where β>1 (*Figure 2b*). In a stable environment, it is advantageous to minimize adaptive timescales (smaller τ_sweep_) and so instances where β<1 will be detrimental. However, in a fluctuating environment, longer adaptive timescales may be advantageous because they avoid a HT selective sweep that may be beneficial in one environment but deleterious in the other. This trade-off between short-term and long-term benefits can be framed in terms of two opposing evolutionary strategies: 1) a low-β strategy with more persistent LT modifications which facilitates rapid environmental tracking with less heritability; and 2) a high-β strategy favoring more rapid selective sweeps of innovative HT modifications at the expense of shorter-term environmental fitness tracking. A low-β strategy could also be viewed as a type of bet-hedging strategy favored under enhanced environmental variability^10^, while a high-β strategy would be favored under stable conditions.

## Evolutionary trade-offs

To illustrate the interaction between evolutionary strategy and realistic environmental fluctuations, we present an example highlighting how the ideal evolutionary strategy for warm temperature adaptation will depend on both physical and biological timescales. Using the output from the global eddy-resolving GFDL Coupled Climate CM2.6 Model^25^ 2xCO2 simulation, we analyzed Lagrangian trajectories released every 2°x2° in the surface (9,200 trajectories per analysis), integrated using the OceanParcels code^26^ (*Methods; Figures S5&S6*). For illustrative purposes, we contrast two populations with environmentally relevant growth rates^27^ – 0.1 day^-1^ (popA) and 1 day^-1^ (popB) – being advected along the same trajectories. We analyzed trajectories for 350 generations for each hypothetical population (2,426 days and 242 days) and calculated environmental fluctuations over each trajectory relative to both a temperature threshold (>28°C) and to the generation time (τ_f_). This highlights how two populations in a single parcel of water can experience the same changes in environmental conditions differently. For example, 30 days in waters >28°C would translate into a τ_f_=4.3 generations for popA and τ_f_= 43 generations for popB. In other words, for the same physical dynamics, a slower growing population (popA) would experience a more variable environment while popB would experience a more stable environment.

Considering both biological timescales, 27-30% of all released particles experienced 28°C at least once within the 350 generation 2xCO2 run. Based on the duration of physical fluctuations (τ_f_) and a conservative estimate of τ_HT_ = 50, we predict that α >1 for 70-79% of the popB trajectories that experienced 28°C. In other words, as a result of a shorter generation time, this population experienced long enough exposure times to >28°C waters that adaption through genetic modifications (HT) should occur. A faster τ_HT_ would increase the fraction with α >1. In contrast, because popA experienced a more variable environment due to its longer generation time, we estimate that selective sweeps (α >1) would occur in only 2-12% of the popA trajectories (*Figure* 3). We confirmed this prediction using 2 representative trajectories (*Supplement S3*). These results suggest that, within a given environment, directional selection is more effective for faster growing marine microbes than slower growing populations, making it more likely for HT selective sweeps to occur. This is because faster growing populations experience the selective environment for a larger number of generations (τ_f_).

Consideration of the β criteria paints a different picture. For the example of warm temperature adaptation, we identified the trajectories where a low-β strategy would be beneficial (*Figure 3*). We find that 41% of the popA trajectories could employ a low-β strategy using reasonable LT transmission timescales (τ_LT_=10-50). This is in contrast to the popB trajectories where only 25% could employ a low-β strategy; 75% of trajectories experienced environmental fluctuations that were either too fast (τ_f_<10) or too slow (τ_f_>50). This analysis highlights an evolutionary trade-off for marine microbes: 1) faster response to variable environments through a low-β strategy where LT modifications provide a competitive advantage versus 2) faster selective sweeps that provide an advantage based on HT modifications. We predict that the bet-hedging low-β strategy with more persistent LT modifications will be favored by organisms that experience subjectively shorter timescale fluctuations. While this example contrasts two populations experiencing the same physical environment, the hypothesis also applies to organisms living in different regions. For example, relatively stable environments (e.g. oligotrophic) should favor a high-β strategy (less LT mechanisms) while more variable environments (e.g. upwelling/coastal) should favor a low-β strategy (more LT mechanisms).

**Figure 3:**
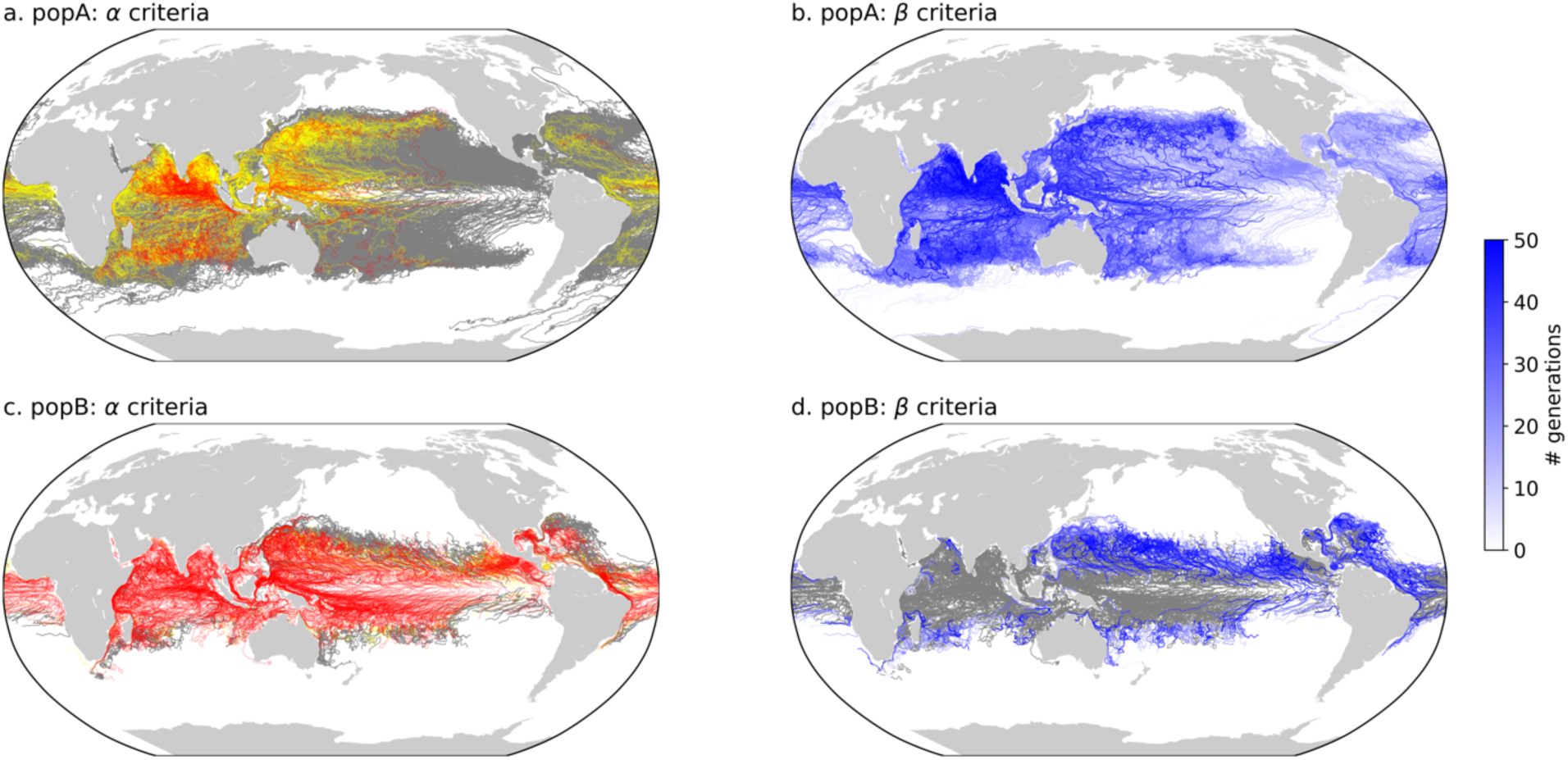
Differences in selective pressure for popA (panels a and b) versus popB (panels c and d). Panels a and c show trajectories predicted to have α>1 and so experience a HT selective sweep. Here we assume that τ_HT_<50 generations and so α>1 for trajectories with mean τ_f_ >50 (red trajectories). This is a conservative estimate since the average model τ_HT_ = 15±7 with max τ_HT_ = 60. Trajectories with the potential for a HT sweep (mean τ_f_ <50 but the maximum τ_f_ >50) are shown in yellow, and trajectories where a sweep is unlikely (maximum τ_f_ <50) are shown in grey. Panels b and d show the estimated timescale of τ_LT_ necessary for a low-β strategy. Trajectories with τ_LT_<50 generations are shown in shades of blue while trajectories with τ_LT_>50 are shown in grey.

Untangling the interactions between the physical timescales of advection and the biological timescales of evolution is necessary to accurately predict how and where marine microbes will adapt to novel environments. Specifically, these results demonstrate that different evolutionary strategies (e.g. low-β versus high-β) are favored by different combinations of fluctuation patterns and cell growth rates and that these strategies can play key roles in shaping microbial fitness and underlying trait values. The importance of the interaction between physical and biological timescales in determining adaptation outcomes brings into question the current assumption of instant acclimation of marine microbes in global carbon cycle models. Understanding these dynamics and constraining marine microbial adaptation timescales will require an improved mechanistic understanding of adaptation that includes variation from LT modifications and the quantification of critical biological timescales including τ_LT_ and τ_HT_. This work suggests that marine microbial populations commonly experience dynamic ocean conditions that favor short-term adaptive strategies (i.e. low-β), and provides a foundation for understanding future shifts in microbial trait distributions and biogeochemical cycling.

## Acknowledgements

This work was supported by grants from the Simons Foundation (509727 and 542389, NML), the Moore Foundation (MMI 7397), NSF (OCE 1538525), NOAA’s Office of Oceanic and Atmospheric Research, and a Royal Society University Research Fellowship to SC. The authors would also like to acknowledge L. Resplandy and J. Busecke for their assistance with the CM2.6 output and Cameron Thrash for comments on the manuscript.

## Author Contributions

NML and NW designed the study, developed the model, analyzed the data, and wrote the manuscript. EZ conducted the trajectory analyses and assisted with writing the manuscript, SC contributed to the experimental design and manuscript, and JD contributed the CM2.6 model simulations and to the writing of the manuscript.

