## Supplementary material for "Hitting a moving target: Microbial evolutionary strategies in a dynamic ocean": Methods and Supplemental Information

#### Methods:

We modeled an individual based adaptive walk using a modified version of Fisher's[1] geometric adaptation model from *Kronholm and Collins*[2] – the EpiGen model. Fitness changes were driven by both LT modifications and HT modifications, where HT modifications were fixed and LT modifications reverted with probability $\mu_{rev}$ (LT reversion rate). Details of the model formulation, model optimization, and simulations are presented below. We then combined the EpiGen model with output from an eddy-resolving climate model (GFDL CM2.6)[3]. The model simulations and methods for the trajectory analysis are described below.

***EpiGen Model formulation:*** Phenotypic space was represented as an *n*-dimensional hypersphere where an individual's phenotype, *z*, was characterized by its distance from the hypersphere origin with radius *r*. Fitness for each individual (*w*) was calculated as:

$$w(z) = e^{(-z^\wedge 2)/2} \quad \text{Eq. 1}$$

such that an individual located at the origin had an optimal fitness of 1 and fitness declined as a Gaussian function as the phenotype moved away from the origin of the hypersphere [4].

The simulations began with $z = r = 1$ for all individuals in the population such that $w$ = 0.6065. The phenotype was altered through both LT and HT modifications which were represented as mutational vectors with random directions and magnitudes in phenotypic space. A new phenotypic value, $Z_{mut}$, was then calculated as[5]:

$$z^2_{mut} = z^2 + m^2 + 2mz\sin(\Theta) \quad \text{Eq. 2}$$

where *m* is the length of the mutational vector and Θ is the angle between the mutational vector and the vector running from the current phenotype to the origin ($\Theta \in [-\pi/2, \pi/2]$). For each new modification, Θ in *n*-dimensional space was drawn from the probability density (*P*) [5]:

$$P = Z\cos^{n-2}(\Theta) \qquad \text{Eq. 3}$$

where $Z$ is a scaling constant and was calculated as:

$$\int_{-\pi/2}^{\pi/2} Z\cos^{n-2}(\Theta)d\Theta = 1 \qquad \text{Eq. 4}$$

All model parameters are given in *Supplemental Table 1*.

The model was initialized with a population of $N$ uniform individuals: here N was varied from $N = 10^3$ to $N = 10^5$. HT modifications ($N_{HT}$ =10) and LT modifications ($N_{LT}$ =90) were then introduced into the population. The modification supply (population size x modification rate) remained constant in each generation and no more than one LT and one HT modification was allowed to occur in a single individual. Eq. 2 was used to calculate new mutant phenotypes. Isotropic modifications in phenotypic space were represented through the uniform distribution of HT modifications, $m_{ht}$, between 0 and $2r$, $m_{ht} \sim U(0, 2r)$, which generated non-uniform fitness effects [2,5]. While LT modifications ($m_{lt}$) were introduced in the same manner as HT modifications, the effects of LT modifications were smaller than HT modifications with a uniform distribution of $m_e$ between 0 and $l$. Hence, $m_{lt} \sim U(0, l)$ instead of $m_{ht} \sim U(0, 2r)$, where $2r$ is the maximum effect of HT modifications and $l \leq 2r$ is the maximum effect of LT modifications. Fitness for each mutant phenotype was then calculated using Eq. 1, and the next generation was then created by sampling from the current population with replacement.

***Simulations:*** We tested the impact of variable selection pressures by introducing intervals in an adaptive walk where the population moved between a 'new environment' (*Figure 1 white shading*) and the 'ancestral environment' (*Figure 1 grey shading*). In the 'new environment', selection was based on fitness in the 'new environment' so the sampling probability of an individual was weighted by its fitness until $N$ offspring had been produced. Selection in the 'ancestral environment' occurred through the stochastic removal of organisms with relatively more HT modifications (i.e. higher HT modification abundance), which corresponds to stabilizing selection. We assume that all modifications have an equal chance of being conditionally deleterious (being neutral or adaptive in the 'selection' or 'new environment', but deleterious in some other environment) so that individuals who have accumulated a high number of modifications in the selection environment have a higher probability of decreased fitness in the ancestral environment.

We tested the impact of the strength of stabilizing selection by running the model in 3 modes for the 'ancestral environment': strong stabilizing selection (strong SS), weak stabilizing selection (weak SS), and neutral selection (Control). In the strong SS regime, selection in the 'ancestral environment' was weighted based on the reciprocal number of HT modifications ($1/N_{HT}$). This resulted in a strong selection against modification load: for example, there would be a 44% selection differential between an organism with 15 HT modifications and one with 2 modifications ($1/2 - 1/15 = 0.44$). Under weak SS, the selection differential was reduced to a maximum of 10% between any given individual. A Control simulation was also conducted where individuals were randomly drawn from the population in the 'ancestral environment' (i.e. neutral selection). The time spent in each environment measured in generations (i.e. interval length ($\tau_f$)) was systematically varied from 10 to 100 generations corresponding to environmental fluctuations of days to several months. This encompasses environmental variability driven by mesoscale eddies to fluctuations driven by advection throughout the global oceans [6].

The LT transmission timescale ($\tau_{LT}$) was varied from no LT modifications ($\tau_{LT}$=1) to maternal effects ($\tau_{LT}$=4 generations) to $\tau_{LT}$=10, 20, 60, and a proof-of-concept long lasting LT effect ($\tau_{LT}$=150 generations). Finally, population size was also varied from (N= $10^3 - 10^5$). $\tau_{HT}$ was an emergent property of the model and varied as a function of mutational supply (a function of population size) and $\tau_f$ which impacts modification effect due to stabilizing selection in the 'ancestral' environment (*Supplementary Figure S1*). Simulations were conducted for 10,000 generations (except when using output from the GFDL Coupled Climate CM2.6 Model, described below) and each simulation was done with 50 replicates. Example output for the 3 model modes are shown in *Supplementary Figure S2.* Model output was analyzed to determine the effects of interval length and selection strength on population fitness. The overall patterns remained unchanged across all sensitivity tests and are discussed in the main text and below (*Supplement S1 & S2, Supplementary Figures S3 & S4*).

While population size does play an important role in adaptive timescales[7], *Kronholm and Collins*[2] demonstrated that the modification supply (population size x modification rate) of largely non-neutral modifications in the model framework is high enough that selection overwhelms drift. This was confirmed with our tests which showed consistent results (i.e. selective HT sweeps) despite significantly increasing population size (*discussed in Supplement S2*).

***Global Trajectory Analysis:***

Lagrangian trajectories were computed with surface velocity and sea surface temperature output from the eddy resolving, 0.1° x 0.1° horizontal resolution, GFDL Coupled Climate CM2.6 Model[3] with 2x$CO_2$ forcing. The 2x$CO_2$ model simulation branched from the control run (pre-industrial CO2) after year 121 and atmospheric $CO_2$ was increased by 1% per year until the concentration was double after which atmospheric $CO_2$ was held constant. The SST for the 2x$CO_2$ simulation is shown in *Supplementary Figure S5*. For this study, we analyzed trajectories initialized on a 2°x2° horizontal grid from 80°S to 70°N (resulting in 9218 trajectories released). Trajectories were integrated using OceanParcels code [8] version 1.0.3 with a timestep of 10 minutes. Location and temperature along the trajectories were recorded for illustration once per day. Two trajectory lengths were analyzed: 2,426 days (6.6 years) and 242 days of output both starting 60 years after the branch. 2,426 days corresponds to 350 generations of a phytoplankton population growing at an average rate of 0.1 $d^{-1}$ and 242 days corresponds to 350 generations of a phytoplankton population growing at an average rate of 1 $d^{-1}$. These growth rates were chosen for illustrative purposes as representative of typical growth rates for eukaryotic phytoplankton[9]. We also analyzed the same trajectories in the control simulation (N=9218) to determine the mean difference in temperature experienced by the particles between the control and 2x$CO_2$ simulation. The average temperature experienced by the particles increased from 16.3 ± 0.8°C in the control simulation (pre-industrial $CO_2$ concentrations) to 17.6 ± 0.9°C in the 2x$CO_2$ run and the fractional area experiencing ≥28°C increased by >200%.

To test the representativeness of the trajectories released on the 2°x2° grid, we analyzed an additional set of trajectories (N= 21,912) that were initialized between 40°S and 40°N on a grid slightly off-set from the original grid (i.e., 2°x2°, 2.1°x2°, 2°x2.1°, and 2.1°x2.1°). The statistics of the intervals above 28°C experienced by the Lagrangian trajectories were calculated for these ensemble runs and shown in *Supplementary Figure S6*. The overall patterns were consistent between the 4 sets of trajectories suggesting that the trajectories on the 2°x2° grid are representative of the model dynamics.

***S1: Adaptation under variable selection and selection strength***

We tested the impact of the strength of stabilizing selection by running the model in three modes (*see Methods*): strong stabilizing selection (strong SS), weak stabilizing selection (weak SS) and a Control. When $\alpha<1$, selective HT sweeps were not observed in either the strong SS or the weak SS simulations and overall population fitness in the new environment was reduced relative to the Control (e.g. *Supplementary Figure S2a*). When $\alpha>1$, selective HT sweeps occurred in both the strong SS and weak SS simulations but HT selective sweeps were never observed in the Control simulations. When $\beta<1$ and selection against HT modification abundance was weak (weak SS), oscillating fitness patterns were observed due to reduced selection differentials between individuals (*Supplementary Figure S2, right column*). Due to this reduced selection differential (e.g. reduced HT diversity), the population as a whole consistently tracked environmental fluctuations using LT modifications both before and after a sweep. Whereas under strong selection, fitness was highly variable and sporadic due to greater intra-population HT differentials between individuals. This enhanced gradient resulted in individuals containing highly beneficial HT modifications being preferentially selected and results in a greater probability of selective sweeps during shorter timescale fluctuations when $\beta<1$ than in the weak SS simulations (*Figure 2; Supplementary Figures S3, S4*). For example, by $\tau_f = 40$ generations with $\beta<1$, 100% of replicates under strong SS exhibited a HT selective sweep whereas only half exhibited sweeps under weak SS (*Supplementary Figure S4*). This suggests that populations containing more closely related individuals (e.g. lower HT diversity) may be slower to adapt as a whole (conservative bet-hedging) than those with higher HT diversity (e.g. increased selection differentials) which may lead to a greater rate of strategy diversification among subpopulations (diversification bet- hedging) [10].

After the initial sweep, mean fitness remained significantly higher (two-sample t-test; $p \ll 0.01$) and less variable (two-sample f-test $p \ll 0.01$) than the fitness prior to the sweep under both weak and strong SS. In summary, certain fluctuating environmental conditions that yield weak selection gradients (i.e. reduced selection strength -- weak SS) can further delay HT-driven adaptation and promote the use of LT modifications for environmental fitness tracking (i.e. low $\beta$-strategy). Alternatively, environmental fluctuations that yield stronger selection gradients (i.e. strong SS) may increase the probability of HT-selective sweeps. This is consistent with empirical observations from microbial systems [11,12].

HT-selective sweeps in the model exhibited a step function increase in fitness while LT-driven short-term fitness changes prior to and immediately following the HT-selective sweeps were highly correlated with normalized distance traveled by LT modifications (LT distance traveled/total distance traveled by both HT and LT modifications[2]) ($R^2 > 0.9$, $p << 0.01$, *main text Fig. 1, Supplementary Figure S2*). This suggests populations can initially experience short-term adaptation due to LT-driven dynamics (where LT modifications are unlikely to be stable over long timescales), as seen in other systems [7,13,14] followed by longer-term stabilization of HT-driven adaptation [13].

***S2: Adaptation under variable population sizes***

We varied population sizes ($N = 10^3$, $10^4$, $10^5$) at two $\tau_f$ values (40 and 100) and 5 $\tau_{LT}$ values under strong SS. HT selective sweeps were observed across all population sizes, $\tau_f$ values, and $\tau_{LT}$ values with $\tau_{HT}$ being negatively correlated with population size (i.e. modification supply). We calculated the percent of replicates that swept within the first 1000 generations for all simulations. For $\tau_f = 40$, the probability of an HT sweep within the first 1000 generations was greater at lower population sizes due to the smaller number of individuals that the HT modification has to sweep through (*Supplementary Figure S7*). However, at $\tau_f = 100$, all replicates swept within the first 1000 generations for all population sizes owing to the increased time of the selection period. Hence, during more rapid environmental fluctuations, the rate of adaptation may be enhanced at lower population sizes if the modification supply (population size x modification rate) of largely non-neutral modifications is high enough that selection overwhelms drift. However, at sufficiently large selection intervals (e.g. $\tau_f = 100$), beneficial modifications tend to always sweep independent of population size. Overall, the results described above were not impacted by population size and the $\alpha$ and $\beta$ criteria still applied.

***S3: Adaptive dynamics in Lagrangian trajectories***

We used the output of 2 representative particle trajectories (P1 and P2) as input into the EpiGen model for both popA (growth rate = 0.1 day$^{-}$1) and popB (growth rate = 1 day$^{-1}$). Adaptation of both populations were analyzed over 350 generations – 2,426 days and 242 days, respectively – and 50 replicate runs were conducted for each population and each trajectory. We selected two growth rates as they are an order of magnitude different and are fairly representative of eukaryotic phytoplankton growth rates[9]. This analysis also assumes that the populations are well adapted to the ancestral environment and so growing at reasonable rates within that environment.

The slower growing microbial population (popA) crossed the 28°C threshold several times within 350 generations with an average interval length above 28°C of 8 ± 11 generations for P1 and 8 ± 13 generations for P2 (i.e. we would predict that $\alpha<1$ unless $\tau_{HT}$ is very fast). This resulted in only 6% and 10% exhibiting an HT selective sweep in P1 and P2, respectively. Conversely, the faster growing microbial population (popB) had an average interval length above 28°C of 107 ± 117 generations for P1 and 58 ± 93 generations for P2 (i.e. we would predict that $\alpha>1$ unless $\tau_{HT}$ is very slow). This resulted in 100% and 98% of the replicates exhibiting an HT selective sweep in P1 and P2, respectively.

These example trajectories with realistic temperature fluctuations confirmed the results of the idealized simulations. In both cases, fitness increases were generally encoded by LT modifications when exposure times to new environments were short, and fixation of genetic mutations (selective HT sweep) happened when directional selection persisted for longer periods. This suggests that adaptation of marine microbes to spatially localized 'novel' environments can occur even if advection through the new environment is relatively rapid. However, the duration of exposure to the novel environment is critical. For example, even when a novel environment is experienced every year, if the interval of exposure is short, the probability of an advantageous genetic mutation fixing is low. Since the fixation time for a beneficial mutation will depend on both the supply of beneficial mutations and the strength of selection, the minimum interval of exposure to novel environments needed for populations to switch from LT to HT adaptation will depend both on the population and the nature of the novel environment. However, the general pattern that we observe of LT-encoded adaptation followed by HT-encoded adaptation will hold as long as both types of adaptation are in principle possible.

### Supplementary Table

**Supplementary Table 1** Parameter values used in the model.

| Parameter | Description | Value |
|---|---|---|
| $N$ | Size of population | 1000 |
| $t$ | Number of generations | 15000 |
| $n$ | Number of hypersphere dimensions | 5 |
| $r$ | Hypersphere radius | 1 |
| $N_{HT}$ | Number of HT modifications | 10 |
| $m$ | Effects of HT modifications | $m_{ht}$ ~ U(0, 2$r$) |
| $N_{LT}$ | Number of LT modifications | 90 |
| $\mu_{rev}$ | LT reversion rate | [0.02, 0.05, 0.15, 0.30, 0.75] |
| $m_e$ | Effects of LT modifications | $m_{lt}$ ~ U(0, $l$) |
| $l$ | Limit for the distribution of $m_e$ for LT effects | 0.3 |

### Supplementary Figures

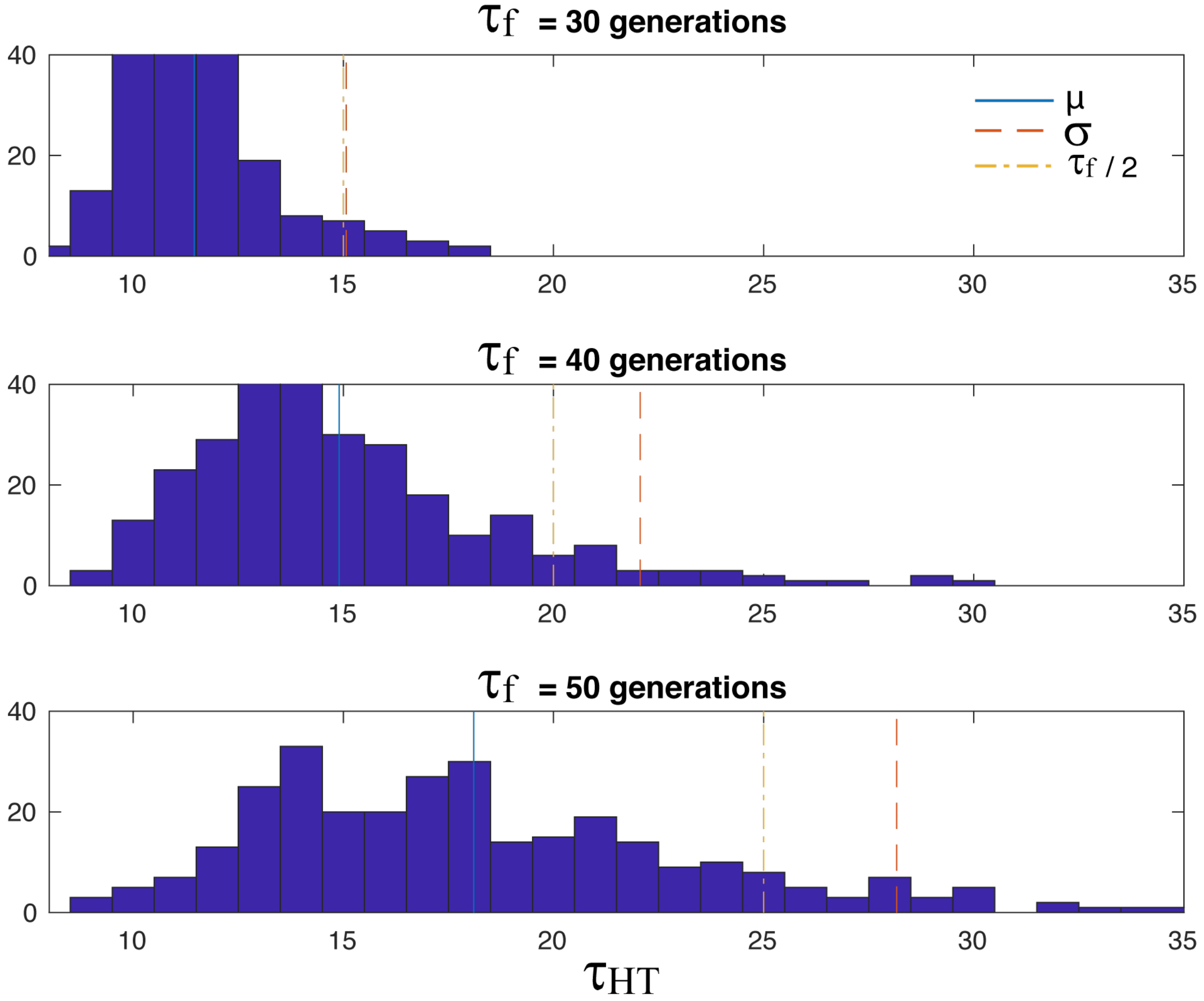


**Figure S1: The distribution of $\tau_{HT}$ as a function of $\tau_f$.** Shown are the distributions and averages of $\tau_{HT}$ as a function of $\tau_f$ which modulates the modification effect (see above). The blue line is the mean $\tau_{HT}$, the red line is the standard deviation, and the yellow line is $\tau_f$ / 2.

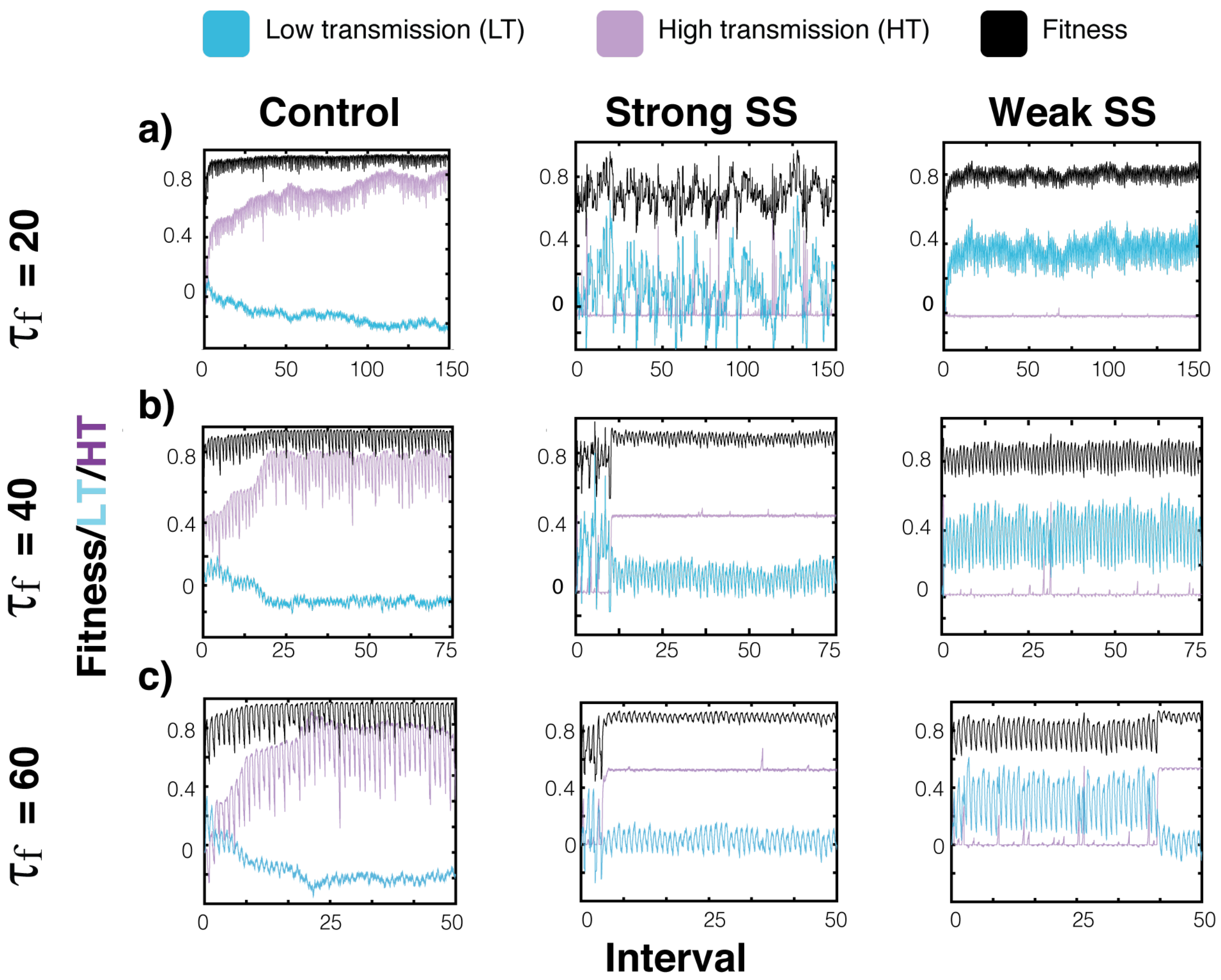


**Figure S2: Fitness, HT, and LT dynamics for Control, Strong SS, and Weak SS runs of select $\tau_f$.** The fitness trajectories (black) for representative simulations are shown concurrently with the underlying relative distance traveled by LT (blue) and HT (purple) changes for $\tau_f$ of 20, 40, and 60 replicates.

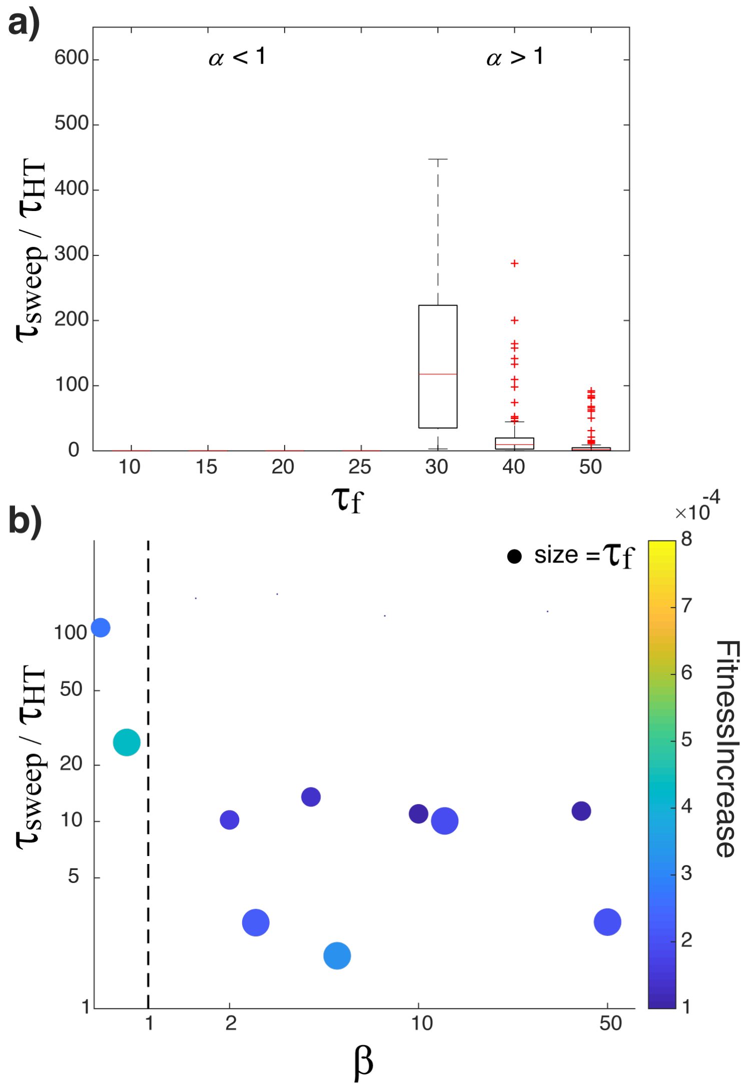


**Fig. S3 Timescales and outcomes of adaptation for weak SS runs determined by the values of α and β.** Panel a illustrates the α criteria by showing the impact of environmental fluctuations ($\tau_f$) on $\tau_{sweep}$ normalized to $\tau_{HT}$. Panel b illustrates the trade-off associated with a low-β strategy by showing the relationship between the rate of fitness increase in a new environment (colorbar) with $\tau_{sweep}$ normalized to $\tau_{HT}$. In panel b, $\tau_f$ is represented by the size of the symbol.

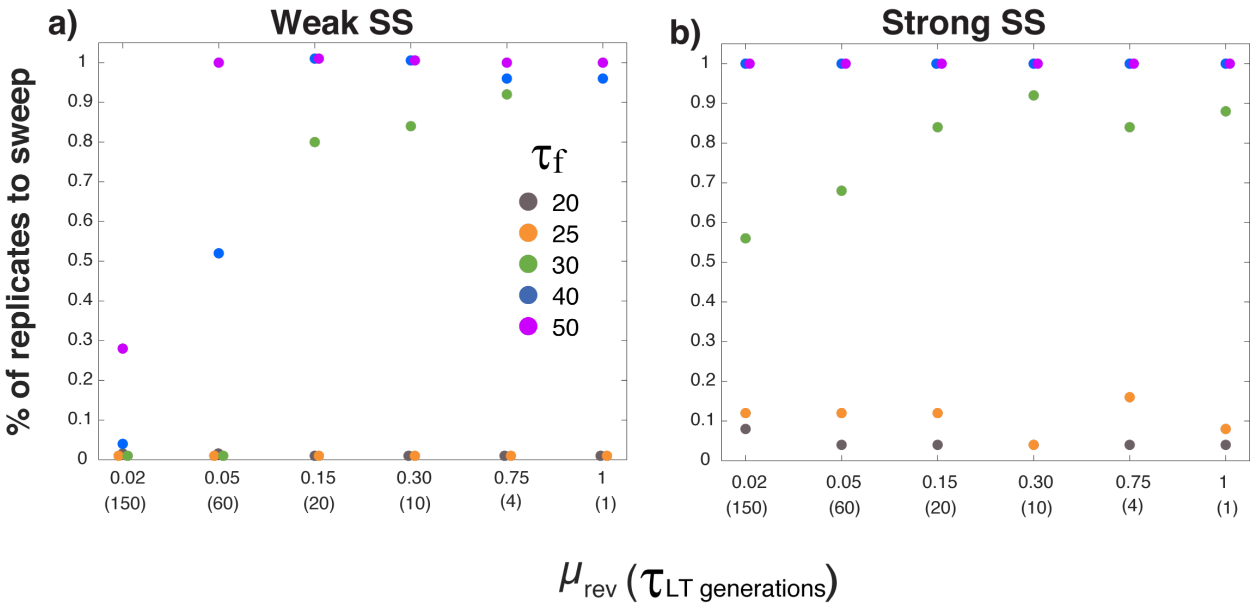


**Figure S4: Percent replicates to sweep for weak SS and strong SS runs as a function of the LT reversion rate $\mu_{rev}$ ($\tau_{LT}$) and $\tau_f$.** Plot a) shows the percent out of 50 replicates (per run type) to sweep for weak SS runs across $\mu_{rev}$ ($\tau_{LT}$) at different $\tau_f$ intervals with each color corresponding to a different $\tau_f$. Plot b) shows the same dynamics as in plot a) but for the strong SS runs.

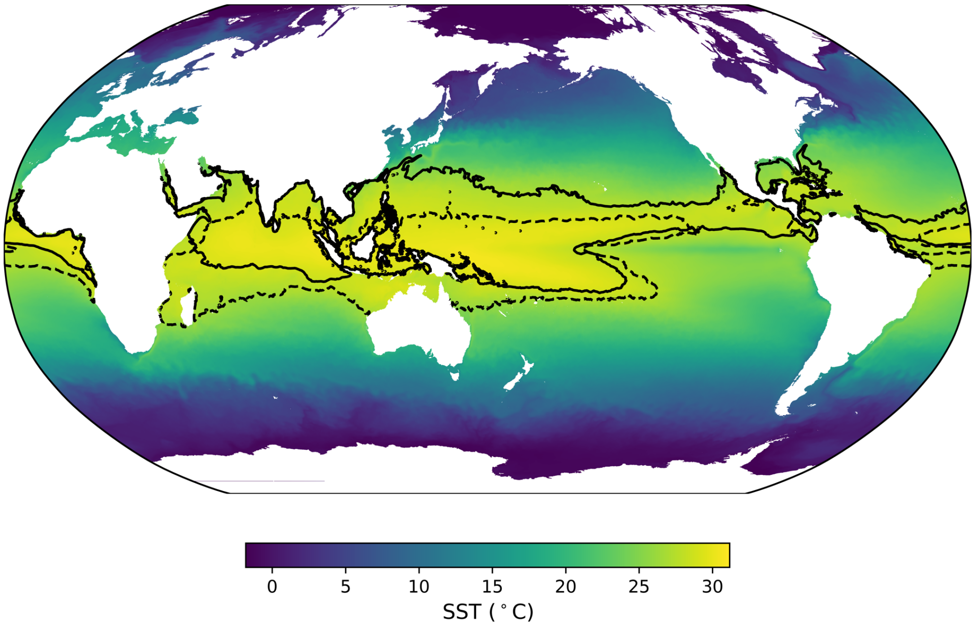


**Figure S5: Annual average temperature** of model year 181 (the first 2xCO2 model year analyzed for this paper) overlaid with 28°C contours: summer average (solid black lines; average of June, July, and August) and winter average (dashed black lines; average of December, January, and February).

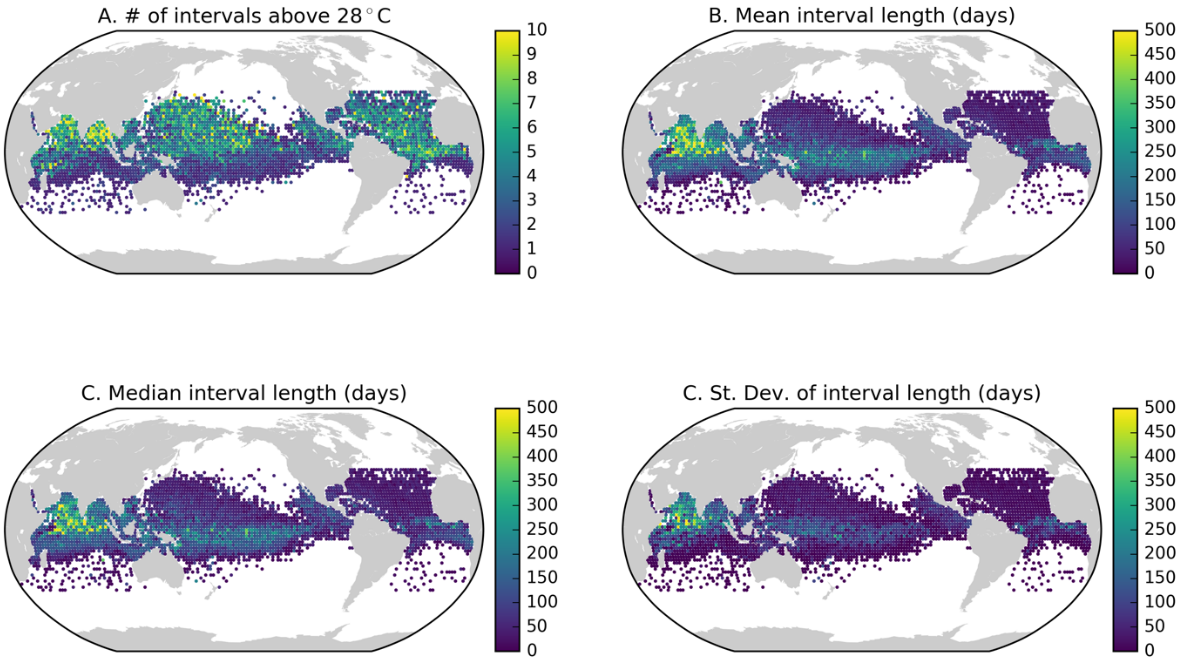


**Figure S6: Statistics of the intervals above 28°C experienced by the Lagrangian trajectories.** Statistics are calculated as the average of four trajectories near each initialized location from 40°S to 40°N (i.e., 2°x2°, 2.1°x2°, 2°x2.1°, and 2.1°x2.1°), and plotted at the location of initialization.

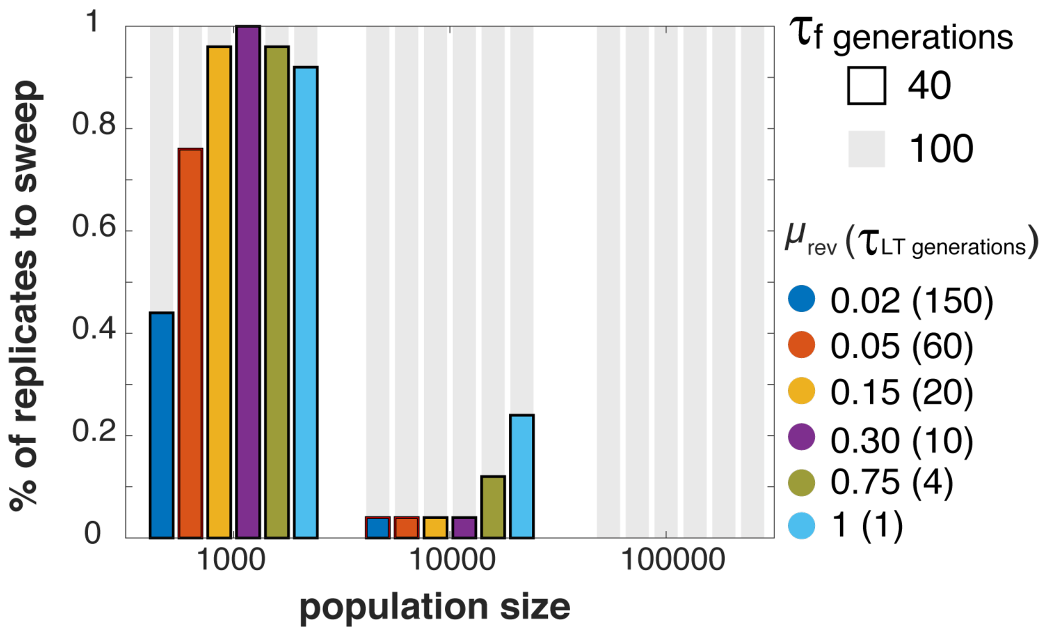


**Figure S7: Percent of replicates to sweep within 1000 generations as a function of the population size and the LT reversion rate $\mu_{rev}$ ($\tau_{LT}$) at $\tau_f = 40$ and $\tau_f = 100$ generations.** Strong SS runs were conducted at $\tau_f = 40$ and $\tau_f = 100$ for 3 population sizes of N = 1000, 10000, 100000. For $\tau_f = 40$ runs, bars are outlined in black and each bar color represents a different $\mu_{rev}$ ($\tau_{LT}$). The percent of replicates (50 reps per run type) to sweep within the first 1000 generations of runs is enhanced at lower population sizes across $\mu_{rev}$ ($\tau_{LT}$) rates. For $\tau_f = 100$ runs, bars are colored in grey and show 100% of replicates sweeping within the first 1000 generations across all $\mu_{rev}$ ($\tau_{LT}$) and all population sizes.
